# A disengaging property of dopamine signaling

**DOI:** 10.1101/2023.11.29.569116

**Authors:** Milan D. Valyear, Noémie M-L. Eustachon, Madeleine M. Morris, Irina Alymova, Bryanna N. Tremblay, Nastasia M. Mitrikeski, Jonathan P. Britt

**Author notes:** Jonathan P. Britt, Department of Psychology, McGill University, 1205 Ave. Dr. Penfield, Montréal, QC, H3A 1B1, Canada.

## Abstract

Dopamine regulates the frequency of motor actions, but it is unclear whether it similarly regulates the duration of discrete behaviors. Here, we show that dopamine signaling within the medial accumbens shell reinforces actions while simultaneously constraining their duration. When mice hold down a lever to continuously self-stimulate dopamine axons in the medial accumbens shell, hold-downs are rarely longer than 5 seconds, and higher stimulation frequencies elicit shorter, but more numerous, lever hold-downs. This disengaging property of medial accumbens shell dopamine also applies to consummatory behavior, as mice are quicker to both terminate and reinitiate bouts of licking paired with high frequency dopamine axon stimulation. Dopamine D1 receptors underpin this disengaging property of dopamine signaling, as we demonstrate a D1 agonist shortens both self-stimulation hold-downs and bouts of consummatory behavior. Altogether, we uncover an anatomically defined role of dopamine, which is to disengage its own reinforcing properties through D1 receptor signaling.

## Introduction

The discovery that rodents^1^ and people^2,3^ perform operant responses to electrically stimulate the medial forebrain bundle ignited speculation about the psychological consequences of this stimulus. Did it elicit feelings of pleasure or substitute for natural goals^4^? In the ensuing search for the neural substrates recruited by this stimulation researchers were quick to appreciate that multiple psychological processes were involved. Rodents^5–7^ and people^2^ prefer brief stimulation trains to continuous ones, and animals are quick to *turn off* brain stimulation even in situations where they are unable to turn it back on^8^. The reinforcing and disengaging properties of medial forebrain bundle stimulation were presumed to arise from distinct neural substrates indiscriminately recruited by the non-specific nature of electrical stimulation^9^.

The reinforcing properties of electrical brain stimulation are well-explained by dopamine signaling in the nucleus accumbens^10^. Dopamine levels in the accumbens, but not dorsal striatum, increase when mice self-stimulate the medial forebrain bundle^11,12^ and pharmacologically elevating dopamine in the medial accumbens shell, but not core, amplifies the reinforcing efficacy of this electrical stimulation^13^. Intriguingly, dopamine release in the medial accumbens shell is not solely implicated in reinforcement. Mice avoid stimulation of accumbens inputs that corelease dopamine and glutamate if glutamate synthesis is blocked in this pathway^14,15^. Additionally, dopamine levels in the medial accumbens shell become elevated in response to appetitive and aversive stimuli^16,17^, unlike in the lateral accumbens shell where dopamine levels dip in response to aversive stimuli^17^. The involvement of dopamine signaling in the medial accumbens shell in both appetitive and aversive processes distinguishes this area from other striatal subregions and raises the possibility that dopamine in this area simultaneously controls behavioral reinforcement and disengagement, perhaps serving to limit the duration of highly reinforced actions^18^.

Animals engage with natural reinforcers by performing discrete bouts of behavior, such as bouts of licking when thirsty^19^. The frequency of engaging in licking behavior reflects complex interactions of taste, homeostatic need, satiety, and learned post-ingestive consequences. However, lick bout durations are primarily driven by taste^19,20^. For example, sweetening a solution with sucrose reliably increases lick bout durations even if it simultaneously reduces total consumption^21^. Thus, the frequency of engaging in a rewarded behavior and the mean duration of engagement reflect overlapping but separable processes with disengagement depending primarily on the sensory properties of the reinforcer. It is unclear what sensory properties, if any, are elicited by dopamine neuron stimulation and whether animals actively regulate the duration of dopamine neuron self-stimulation like they regulate the duration of engagement with natural reinforcers. However, rodents can learn to control the duration of intravenous cocaine infusions, which directly elevate striatal dopamine levels^22,23^.

The downstream substrates recruited by electrical medial forebrain bundle stimulation and taste stimuli overlap^24^, but their exact identities remain obscure. A recent report suggests that strong, but not weak, stimulation of ventral tegmental area dopamine neurons, a consequence of electrical medial forebrain bundle stimulation^25^, evokes a rich and discernable sensory experience^26^. The sensory features of elevated dopamine signaling are speculative, but it is curious that direct excitation of ventral tegmental area dopamine neurons^27^ and downstream D1 receptor-expressing neurons in the medial accumbens shell both halt ongoing bouts of licking^28^ which is broadly consistent with a role in disengaging behavior.

Here, we hypothesize that dopamine signaling in the medial accumbens shell simultaneously reinforces behavior while promoting behavioral disengagement, possibly to prevent the perseveration of highly reinforced actions. To test this hypothesis, we gave mice access to a lever during 30 s trials that when held down provided continuous photostimulation of dopamine axons in the medial accumbens shell. The frequency of this stimulation changed randomly between 2.5, 10, and 40 Hz at the start of each trial. We found that mice gradually increased the number and duration of lever hold-downs until they were cumulatively holding down the lever for a third of the time it was available. After extended training, mice learned to rapidly adjust their self-stimulation behavior in response to trial-by-trial changes in the stimulation frequency. Stimulation at 40 Hz supported the most numerous but briefest lever hold-downs, suggesting that medial accumbens shell dopamine is both reinforcing and disengaging. Modulating dopamine neurotransmission with a D1 receptor agonist shortened, whereas a D2 receptor antagonist lengthened, the duration of lever hold-downs at all stimulation frequencies. Separately, dopamine axon stimulation and a D1 receptor agonist both reduced lick bout durations during consumption of a natural reinforcer. Thus, we report a disengaging property of medial accumbens shell dopamine that is supported by neurotransmission at D1 receptors, which we propose serves to prevent behavioral perseveration by limiting the duration of highly reinforced actions.

## Results

### Mice rapidly disengage high frequency dopamine axon stimulation

DAT-Cre mice were trained to hold-down an active lever to optogenetically self-stimulate dopamine axons in the medial accumbens shell (**Fig 1a-d**). The frequency of stimulation available to the mouse varied randomly between 2.5, 10, or 40 Hz at the start of each 30 s trial (48 trials/session). In selecting these stimulation frequencies, we considered the amplitude of optically-evoked dopamine release in the medial accumbens shell across a range of different frequencies and durations of photostimulation in a separate cohort of mice. We found that 40 Hz photostimulation elicited significantly more dopamine release than 10 Hz photostimulation [*t*_(6)_=9.93, *p*<.001; *F*_(4, 24)_=54.57, *p*<.001; **Fig 1e**], and the amount of release positively scaled with the duration of stimulation [*F*_(4, 24)_=6.75, *p*<.001; **Fig 1f**]. Thus, the amplitude of optically-evoked dopamine release, as measured with rGRAB-DA1m fluorescence in the medial accumbens shell, is a function of both the frequency and duration of photostimulation.

**Figure 1.**
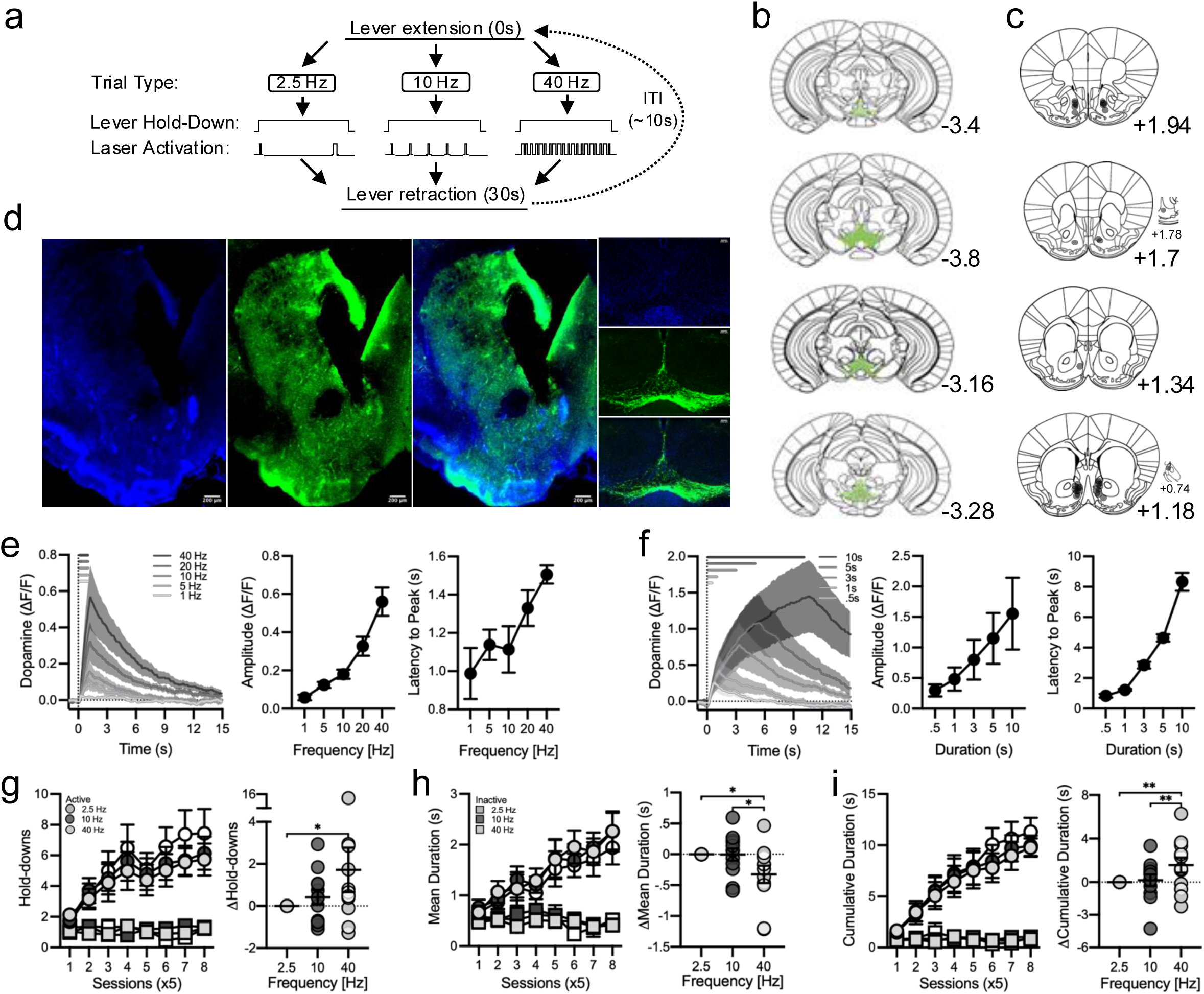
Mice rapidly disengage high frequency dopamine axon stimulation. **a**, Active and inactive levers were inserted into the conditioning chamber for 30 s trials (48 trials/session). Active lever hold-downs triggered photostimulation of dopamine axons in the medial accumbens shell at 2.5, 10, or 40 Hz, which terminated when the lever was released. This schematic shows a fictitious active lever hold-down lasting 0.5 s at each stimulation frequency. **b**, DAT-Cre mice (n=10f, 10m) received bilateral microinfusions of the viral construct AAV5-EF1a-DIO-hChR2(H134R)-eYFP in the ventral tegmental area and **c**, implantation of optical fibres in the medial accumbens shell, which are shown on modified panels from the atlas of Franklin and Watson (2007). Representative images (5x) showing viral expression (eYFP, green) and nuclei (DAPI, blue) in the **d**, striatum (left) with a fibre tract terminating in the medial accumbens shell and ventral tegmental area (right). **e**, rGRAB-DA1m fluorescence (n=4f, 3m) indicative of dopamine release in the medial accumbens shell evoked by 1 s optical stimulation of ventral tegmental area dopamine neurons at 1, 5, 10, 20, and 40 Hz (5 ms pulse; left). The amplitude (middle) of evoked dopamine release increased with stimulation frequency [1 vs 5 Hz *t*_(6)_=1.73, *p*=.97; 5 vs 10 Hz *t*_(6)_=1.47, *p*=.97; all other, *t*_(6)_>3.2, *p*<.038; Frequency, *F*_(4, 24)_=54.57, *p*<.001]. The latency to peak (right) also increased with stimulation frequency but to a much smaller degree [1 vs 40 Hz, *t*_(6)_=3.42, *p=*0.022; all other, *t*_(6)_<2.59, *p>*0.159; Frequency, *F*_(4, 24)_=3.62, *p*=.019]. **f**, rGRAB-DA1m fluorescence evoked by 40 Hz optical stimulation of ventral tegmental area dopamine neurons for .5, 1, 3, 5, and 10 s (5 ms pulse; left). The amplitude (middle) of evoked dopamine release increased with the duration of optical stimulation [.5 vs 5s, *t* _(6)_=3.09, *p*=.049; .5 vs 10s, *t* _(6)_=4.55, *p*=.001; 1 vs 10s, *t* _(6)_=3.88, *p*=.007; all other, *t* _(6)_<2.73, *p*>.116; Frequency, *F*_(4, 24)_=6.75, *p*<.001]. The latency to peak (right) also increased with stimulation duration [.5 vs 1s, *t*_(6)_=.85, *p*=.999; all other, *t* _(6)_>3.59, *p*<.015; Frequency, *F*_(4, 24)_=90.5, *p*<.001]. **g**, Across blocks of 5 training sessions, mice (n=7f, 5m) increased the number of active but not inactive lever hold-downs performed during 30 s trials, particularly during high-frequency trials [left; Lever x Frequency x Session, *F*_(14, 154)_=2.68, *p*=.002]. During the last block of training, mice performed more active hold-downs during 40 Hz trials than 2.5 Hz trials (right). Data in the right panels of *g-i* are shown as within-subject differences from the lowest stimulation frequency (2.5 Hz). **h**, The mean duration of active, but not inactive, lever hold-downs increased over training [left; Lever x Session, *F*_(7, 77)_=4.51, *p*<.001] and active hold-downs were briefest during 40 Hz trials through the last block of training (right). **i**, Mice also increased the cumulative time spent holding down the active but not inactive lever, particularly during high frequency trials, across training [left; Lever x Frequency x Session, *F*_(14, 154)_=2.32, *p*=.006]. The cumulative duration of active lever hold-downs was longest during 40 Hz trials (right). Averaged data are mean ± s.e.m. Symbols represent data from individual mice. Asterisks indicate Bonferroni-corrected and multiplicity-adjusted post-hoc comparisons *p*<.01**, *p*<.05*.

In our lever hold down assay, mice performed more active than inactive lever hold-downs within the first five sessions of training [Lever, *F*_(1, 22)_=7.12, *p*<.022; **Fig 1g**], confirming a reinforcing property of dopamine axon photostimulation in the medial accumbens shell. In these early training sessions, self-stimulation behavior did not vary with trial-by-trial changes in the stimulation frequency [Lever x Frequency, *F*_(2, 22)_=.95, *p*=.40; **Fig S1a-f**], which suggests the immediate sensory property of medial accumbens shell dopamine, while reinforcing, did not differ enough between stimulation frequencies to support rapid adjustments in behavior. With extended training, however, mice came to update their behavior in response to trial-by-trial changes in stimulation frequency, as they exhibited higher rates of lever pressing specifically during 40 Hz stimulation trials [Lever x Session, *F*_(7, 77)_=8.54, *p*<.001; Lever x Frequency x Session, *F*_(14, 154)_=2.68, *p*=.002; **Fig 1g**]. Thus, mice learned to rapidly adjust their self-stimulation behavior within 30 s trials, and the reinforcing property of medial accumbens shell dopamine was most pronounced at higher stimulation frequencies.

To better examine how stimulation frequency influenced self-stimulation behavior trial-by-trial, we calculated the absolute number, mean duration, and cumulative duration of lever hold-downs in the last block of training as a function of trial type (**Fig 1g-i**). All analyses were conducted on absolute measures, but we also show differences in behavior relative to 2.5 Hz trials, the lowest stimulation frequency tested, to decompose significant interactions. At the end of training, mice performed more active lever hold-downs during 40 Hz trials relative to 2.5 Hz trials [*t*_(11)_=2.99, *p*=.02; Lever x Frequency, *F*_(2, 22)_=2.61, *p*=.096; **Fig 1g**], but the mean duration of active lever hold-downs on 40 Hz trials was briefer relative to both 2.5 Hz and 10 Hz trials [2.5 Hz, *t*_(11)_=3.06, *p*=.017; 10 Hz, *t*_(11)_=3.02, *p=*.019; Lever x Frequency, *F*_(2, 22)_=5.3, *p*=.013; **Fig 1h**]. Considering that the stimulation frequency was randomly determined at the start of each trial, these data indicate that mice were rapidly adjusting their self-stimulation behavior in response to changes in the sensory experience of the stimulation. While the highest stimulation frequency was relatively more reinforcing, supporting more active lever hold-downs, mice were fastest to disengage this stimulation, supporting the briefest mean hold-down duration.

The reduced mean hold-down duration at high stimulation frequencies was accompanied by a change in the distribution of hold-down durations. Active hold-downs were seldom longer than ∼5 s at any stimulation frequency (**Fig S2a**), but sub ∼1 s hold-downs were more common during 40 Hz trials. Consequently, the distribution of active hold-down durations was leftward shifted on 40 Hz trials relative to 2.5 and 10 Hz trials [2.5 Hz, D=.15, *p*<.001; 10 Hz, D=.18, *p*<.001; **Fig S2b**]. The small number of active hold-downs longer than ∼5 s suggests an inherent disengaging property of medial accumbens shell dopamine that is amplified at high stimulation frequencies. Together, these data demonstrate that mice will hold down a lever for continuous photostimulation of dopamine axons in a manner that depends on the stimulation frequency, with lower frequencies supporting longer active hold-downs and higher frequencies supporting more numerous active hold-downs. These results are consistent with the hypothesis that high-frequency dopamine activity impinges upon a downstream substrate in the medial accumbens shell that disengages highly reinforced actions.

In the last block of training, when mice were performing more than 200 total active lever hold-downs per session (**Fig S3a**) and cumulatively holding down the active lever for a third of the available trial time (i.e., ∼10 s per trial; **Fig 1i**), cumulative hold-down times were significantly longer during 40 Hz trials [10 Hz, *t*_(11)_=3.37, *p=*.008; 2.5 Hz, *t*_(11)_=3.79, *p*=.003; Lever x Frequency, *F*_(2, 22)_=3.59, *p*=.04; **Fig 1i**], but the absolute difference was small (∼38% versus 33% of total trial time). These active lever presses were clearly supported by dopamine axon photostimulation, as the number, mean duration, and cumulative duration of inactive lever hold-downs did not vary across stimulation frequencies [all, *t*_(11)_<.18, *p*>.99; **Fig S1g-i**]. Moreover, control mice lacking opsins, but with optic fibers targeting the medial accumbens shell, did not distinguish between the two levers or across stimulation frequencies, and performed fewer than ten lever hold-downs during the last week of training (**Fig S3b-e**).

### Dopamine receptor activity regulates the duration of self-stimulation behavior

Dopamine neurons release multiple neurotransmitters in the nucleus accumbens^29,30^, any one of which could promote disengagement. Given recent demonstrations of an aversive component of dopamine signaling in this area^17,31^ and the observation that stimulation of D1 receptor-expressing accumbal neurons terminates ongoing lick bouts^27,28^, we hypothesized that dopamine receptor activity could simultaneously mediate reinforcement and disengagement. To test this idea, we administered (i.p.) agonists and antagonists to D1 and D2 dopamine receptors prior to test sessions using a Latin-square design in the same well-trained mice. The D1 agonist SKF38393 (.25 mg/kg) and D2 agonist quinpirole (.125 mg/kg) were used to augment receptor activity, whereas the D1 antagonist SCH23390 (.02 mg/kg) and the D2 antagonist raclopride (.03 mg/kg) were used to blunt receptor activity. These drug doses were chosen because they have been extensively shown to not affect motor behavior in mice^32–41^. We analyzed drug treatment effects on the absolute number, mean duration, and cumulative duration of lever hold-downs (**Fig 2a,c,e**), but we also show treatment effects normalized to the vehicle condition to decompose significant interactions (**Fig 2b,d,f**).

**Figure 2.**
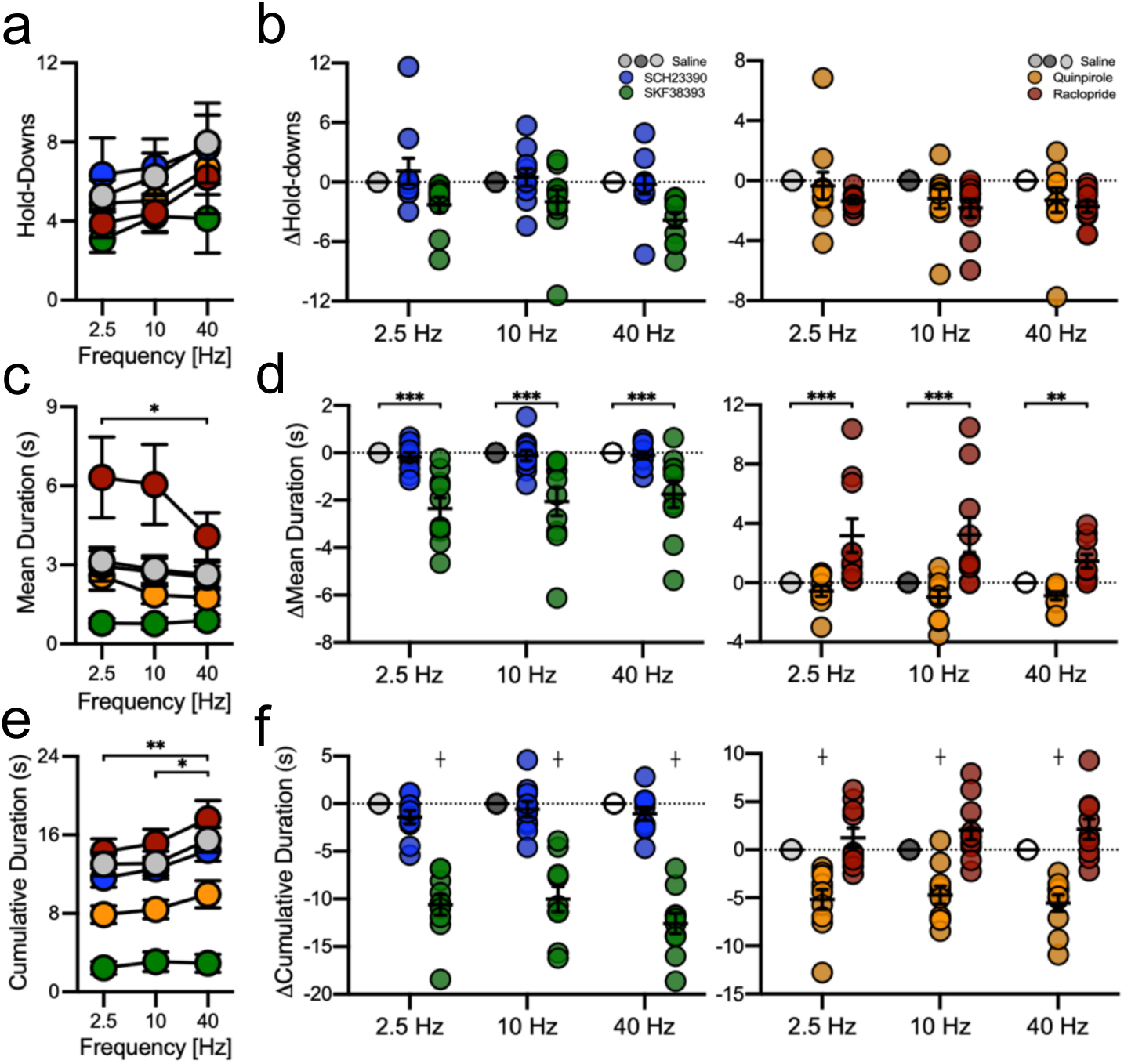
Dopamine receptor activity regulates the duration of self-stimulation behaviour. **a**, The mean number of active lever hold-downs per trial was not affected by administration of the **b**, D1 antagonist SCH23390 or the D1 agonist SKF38393 nor by the D2 agonist Quinpirole or the D2 antagonist raclopride. Data in panels *b*, *d*, and *f* are shown as within-subject differences from the saline treatment condition. **c**, The mean duration of active lever hold-downs was briefer during 40 Hz trials than 2.5 Hz trials (see Fig S4) and modulated in a frequency-dependent manner by both the D1 agonist SKF38393 and the D2 antagonist raclopride. **d**, The D1 agonist SKF38393 shortened whereas the D2 antagonist raclopride lengthened mean hold-down duration relative to saline. The D1 antagonist SCH23390 and D2 agonist quinpirole did not affect mean hold-down duration. **e**, The cumulative duration of active lever hold-downs was longer during 40 Hz trials than 10 or 2.5 Hz trials (see Fig S4). **f**, The D1 agonist SKF38393 and D2 agonist quinpirole reduced the cumulative duration of active hold-downs irrespective of stimulation frequency. Neither the D1 nor D2 antagonist affected cumulative hold-down duration. Averaged data are mean ± s.e.m. Symbols represent data from individual mice. Asterisks indicate Bonferroni-corrected and multiplicity-adjusted post-hoc comparisons *p*<.001***, *p*<.01**, *p*<.05*. Crosses indicate similarly corrected simple effects of treatment relative to saline *p*<.0001┼.

Although dramatic effects on self-stimulation rates have been observed with higher doses of these drugs^42^, the low doses used here did not significantly affect the number of lever hold-downs [Frequency x Treatment, *F*_(8, 72)_=1.47, *p*=.18; **Fig 2a-b, S4a**]. There was, however, a significant drug effect on the mean duration of active hold-downs [Frequency x Treatment, *F*_(8, 72)_=4.95, *p*=.0002; **Fig 2c**]. Relative to saline, the D1 receptor agonist SKF38393 shortened mean hold-down durations, whereas the D2 receptor antagonist raclopride lengthened them [SKF38393: 2.5 Hz, *t*_(9)_=6.22, *p*<.001; 10 Hz, *t*_(9)_=6.04, *p*<.001; 40 Hz, *t*_(9)_=5.13, *p*<.001; Raclopride: 2.5 Hz, *t*_(9)_=11.01, *p*<.001; 10 Hz, *t*_(9)_=12.32, *p*<.001; 40 Hz, *t*_(9)_=6.83, *p*=.0002; **Fig 2d**]. Changes to mean hold-down durations were most pronounced at lower stimulation frequencies, which may reflect a decreased capacity of these drugs to compete with evoked dopamine release at higher stimulation frequencies. Neither the D1 antagonist SCH23390 nor the D2 agonist quinpirole affected the mean duration of active lever hold-downs relative to saline [all, *t*_(9)_<2.84, *p*>.17], consistent with prior work showing that differences in the affinity of dopamine for D1 and D2 receptors results in behavior being more sensitive to increases in D1, and decreases in D2, receptor activity^43^.

The cumulative duration of active lever hold-downs varied across stimulation frequencies [Frequency, *F*_(2, 18)_=11.04, *p*<.001; **Fig S4c**] and drug treatments [Treatment, *F*_(4, 36)_=73.3, *p<*.001], but these factors did not interact [Frequency x Treatment, *F*_(8, 72)_=1.30, *p*=.26; **Fig 2e**]. The D1 agonist SKF38393 and D2 agonist quinpirole both reduced cumulative hold-down durations across all stimulation frequencies [SKF38393: *t*_(9)_=12.94, *p*<.001; Quinpirole: *t*_(9)_=6.01, *p*<.001; **Fig 2f**], whereas the D1 antagonist SCH23390 and the D2 antagonist raclopride did not significantly affect cumulative hold-down durations [all, *t*_(9)_<2.12, *p*>.41]. Inactive lever hold-downs were largely unaffected by drug treatments (see **Fig S5**).

The disengaging property of high frequency dopamine axon stimulation seen during training was observed across all treatment conditions at test [40 vs 2.5 Hz, *t*_(9)_=3.36, *p*=.011; Frequency, *F*_(2, 18)_=5.72, *p*=.012; **Fig 2c**, **S4b**] and was amplified by augmenting neurotransmission at D1 receptors. Together, the relatively brief active lever hold-down durations observed at higher frequencies of dopamine axon stimulation and with a D1 receptor agonist suggest that dopamine signaling at D1 receptors promotes behavioral disengagement and may be a means of preventing the perseveration of highly reinforced actions.

### Stimulation frequency and dopamine receptor activity influence the first lever hold-down

As mice could not predict which stimulation frequency would be available on any given trial, the latency to initiate the first active lever hold-down on each trial did not vary across stimulation frequencies [Frequency, *F*_(2, 18)_=.049, *p*=.95; **Fig S6**]. However, the duration of the first active lever hold-down and the latency to initiate a second were both significantly shorter on 40 Hz trials than 2.5 Hz trials [Duration: *t*_(9)_=3.04, *p*=.021; Frequency, *F*_(2, 18)_=5.01, *p*=.019; **Fig 3a, S7a**; Latency: *t*_(9)_=2.79, *p*=.036; Frequency, *F*_(2, 18)_=4.02, *p*=.036; **Fig 3c, S7b**]. Brevity in the latency to initiate a second hold-down on high-frequency trials is consistent with a reinforcing property dopamine signaling in the medial accumbens shell. The capacity of a D1 agonist and D2 antagonist to modulate the duration of active hold-downs was also present on the first hold-down [Frequency x Treatment, *F*_(8, 72)_=2.61, *p*=.014]. The D1 agonist SKF38393 shortened, whereas the D2 antagonist raclopride lengthened, the duration of the first active lever hold-down relative to saline [SKF38393: 2.5 Hz, *t*_(9)_=8.21, *p*<.001; 10 Hz, *t*_(9)_=6.87, *p*<.001; 40 Hz, *t*_(9)_=6.63 *p*<.001; Raclopride: 2.5 Hz, *t*_(9)_=8.15, *p*<.001; 10 Hz, *t*_(9)_=9.84, *p*<.001; 40 Hz, *t*_(9)_=5.08, *p<*.001; **Fig 3b**]. While the D2 agonist quinpirole did not affect mean hold-down durations overall, it did shorten the duration of these first active lever hold-downs [2.5 Hz, *t*_(9)_=3.86, *p=*.007; 10 Hz, *t*_(9)_=3.61, *p*=0.017; 40 Hz, *t*_(9)_=4.14, *p*=.003; **Fig 3b**]. The latency to initiate a second active hold-down was unaffected by drug treatments [Treatment, *F*_(8, 18)_=.98, *p*=.43; **Fig 3d**]. Thus, on the first active hold-down when mice could not know what stimulation frequency was available, they were quicker to disengage high-frequency stimulation, and this effect was amplified by augmenting neurotransmission at D1 receptors.

**Figure 3.**
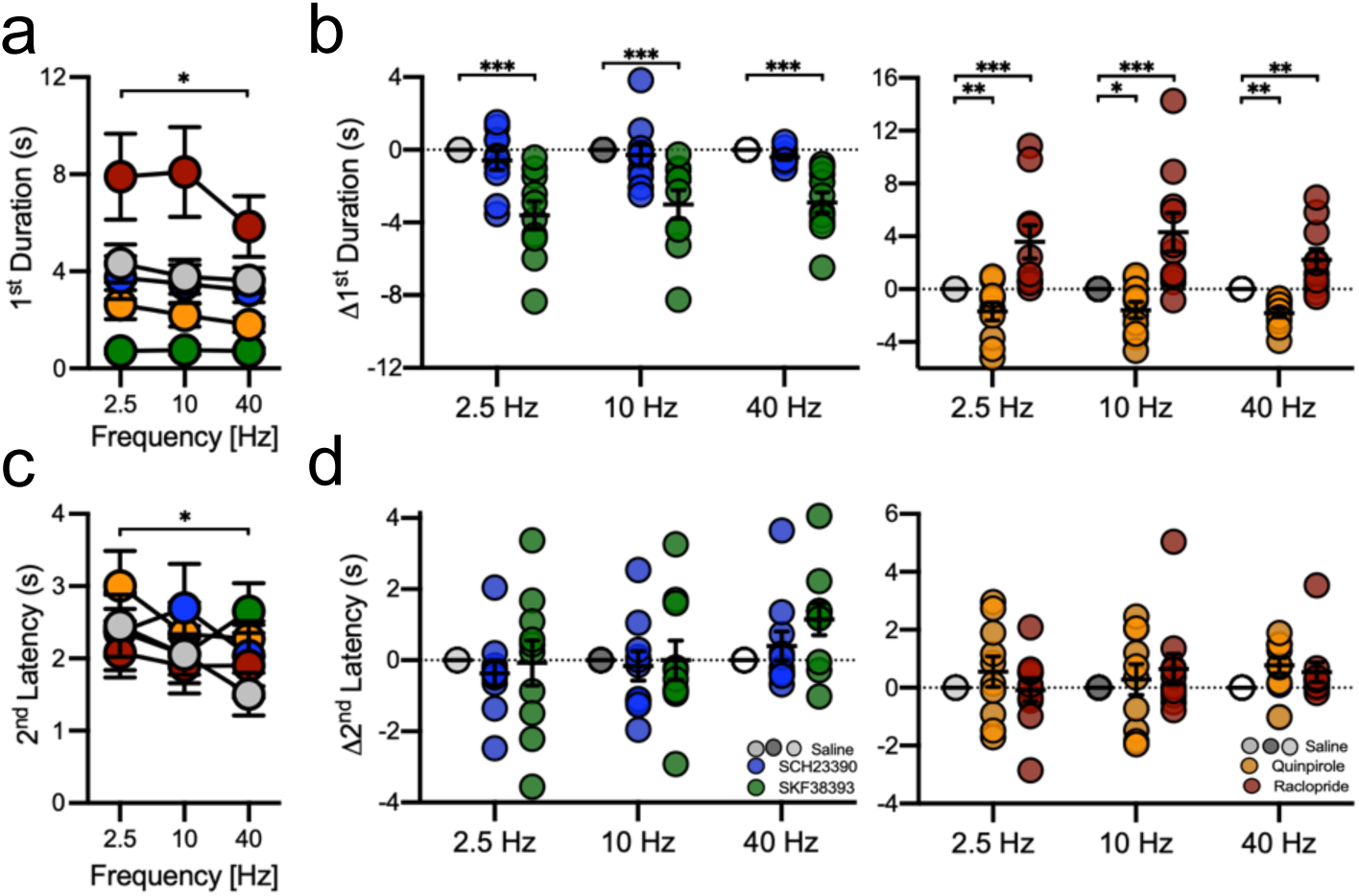
Stimulation frequency and dopamine receptor activity influence the first lever hold-down. **a**, The mean duration of the first active lever hold-down during a trial was briefer during 40 Hz trials than 2.5 Hz trials (see Fig S7) and modulated in a frequency-dependent manner by the D1 agonist SKF38393, D2 agonist quinpirole, and D2 antagonist raclopride. **b**, The D1 agonist SKF38393 and D2 agonist quinpirole both shortened whereas the D2 antagonist raclopride lengthened the mean duration of the 1^st^ hold-down. Data in panels *b* and *d* are shown as within-subject differences from the saline treatment condition. **c**, Mice were quicker to initiate a 2^nd^ active lever hold-down during 40 Hz than 2.5 Hz trials (see Fig S5). **d**, The latency to perform a 2^nd^ hold-down was unaffected by treatment with dopamine receptor agonists and antagonists. Averaged data are mean ± s.e.m. Symbols represent data from individual mice. Asterisks indicate Bonferroni-corrected and multiplicity-adjusted post-hoc comparisons *p*<.001***, *p*<.01**, *p*<.05*.

### Dopamine axon stimulation and D1 receptor activation reduce lick bout durations

We next examined whether increases in medial accumbens shell dopamine had the capacity to disengage naturalistic behavior, specifically food consumption. A naïve cohort of DAT-Cre mice were given unlimited access to a highly palatable sweetened condensed milk solution and received photostimulation of dopamine inputs to the medial accumbens shell at 2.5, 10 and 40 Hz during a random selection of lick bouts. When mice received no stimulation (NS), bouts were ∼3 s long, similar to the duration of self-stimulation hold-downs (**Fig S8a-c**), and all frequencies of dopamine axon stimulation reduced lick bout durations [2.5 Hz, *t*_(7)_=4.73, *p*=.0007; 10 Hz, *t*_(7)_=4.3, *p*=.002; 40 Hz, *t*_(7)_=3.65, *p=*.009; Frequency, *F*_(3, 21)_=9.31, *p*=.0004; **Fig 4a**]. Thus, while palatable foods elevate dopamine levels in the medial accumbens shell^44–47^, augmenting this release by stimulating dopamine inputs, even at low frequencies, hastens the disengagement of consummatory behavior. Still, mice were fastest to initiate a subsequent bout of licking after 40 Hz stimulation-paired lick bouts [2.5 Hz vs 40 Hz, *t*_(7)_=3.26, *p=*.022; NS vs 40 Hz, *t*_(7)_=2.79, *p=*.027 *uncorrected*; Frequency, *F*_(3, 21)_=3.63, *p*=.0297; **Fig 4b**]. Mice concurrently had access to water, and these bouts were seldom, short, and unaffected by dopamine axon simulation (see **Fig S8d-g**). Thus, reinforcing and disengaging properties of medial accumbens shell dopamine are evident during naturalistic consummatory behavior, further emphasizing a role of this substrate in disengaging reinforced actions.

**Figure 4.**
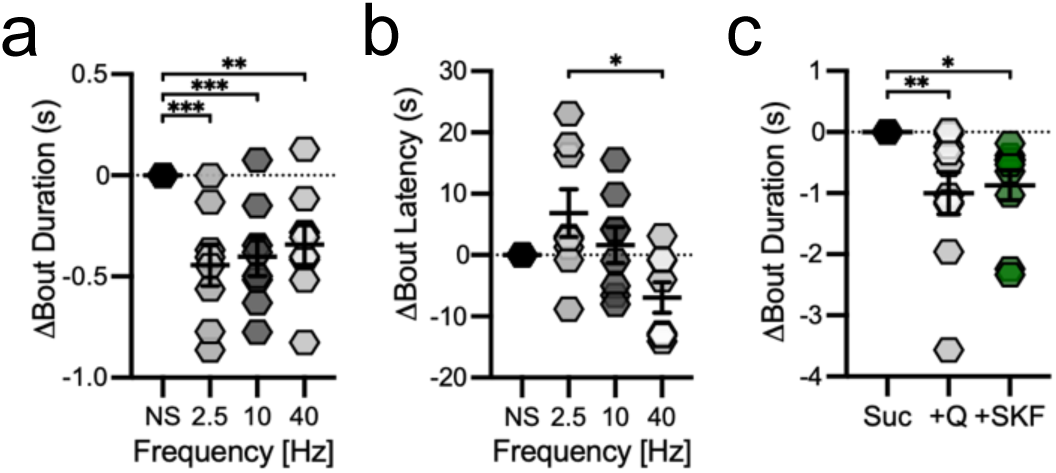
Dopamine axon stimulation and D1 receptor activation reduce lick bout durations. **a,** Mice (n=3f, 5m) shortened the duration of bouts of licking when bouts co-occurred with dopamine axon stimulation at any frequency, but **b**, mice were quickest to initiate a subsequent bout of licking following bouts paired with 40 Hz stimulation. **c**, When an 8% sucrose solution was adulterated with bitter quinine, mice (n=6f, 4m) shortened their lick bouts, and this effect was recapitulated by administering the D1 agonist SKF38393. Data in are shown as within-subject differences from the no stimulation (NS) or vehicle condition. Averaged data are mean ± s.e.m. Symbols represent data from individual mice. Asterisks indicate Bonferroni-corrected and multiplicity-adjusted post-hoc comparisons, *p*<.001***, *p*<.01**, *p*<.05*.

We next sought to determine whether augmenting D1 receptor activity was sufficient to shorten lick bout durations during free consumption of a sweet sucrose solution. We compared the effect of the D1 receptor agonist SKF38393 (i.p.) on lick bout durations to that produced by the adulteration of a sweet solution with bitter quinine, which is well known to reduce lick bout durations. The D1 agonist SKF38393 reduced lick bout durations to the same extent as quinine adulteration [Quinine, *t*_(7)_=3.73, *p*=.0046; SKF38393, *t*_(7)_=3.25, *p*=.0132; Treatment, *F*_(2, 18)_=8.25, *p*=.0029; **Fig 4c**]. Thus, combining naturalistic reinforcing (sweet) and disengaging (bitter) tastes or augmenting D1 neurotransmission similarly disengage consummatory behavior.

## Discussion

Early studies of electrical medial forebrain bundle stimulation implicated the medial accumbens shell as a substrate underpinning its reinforcing properties^12,13^, while other psychological consequences of this stimulation were chalked up to the indiscriminate recruitment of diverse neural substrates near the electrode. Here, by allowing mice to control the duration of dopamine axon stimulation in the medial accumbens shell, we demonstrate that this single substrate has both reinforcing and disengaging properties. Changes in the number and mean duration of lever hold-downs diverge as the frequency of dopamine axon stimulation increases. At higher stimulation frequencies the number and cumulative duration of hold-downs grows, consistent with stronger reinforcement, whereas the mean duration of hold-downs wanes, consistent with disengagement. Moreover, augmenting neurotransmission at D1 receptors shortened, whereas blunting neurotransmission at D2 receptors lengthened, the mean duration of active lever hold-downs, even on the first hold-down. The relatively brief hold-down durations for high-frequency self-stimulation and in response to a D1 receptor agonist identifies dopamine signaling at D1 receptors as a substrate serving to disengage highly reinforced actions, even naturalistic consummatory actions, potentially to prevent behavioral perseveration.

The frequency of dopamine axon stimulation impacted the duration of the first lever hold-down of each trial as well as the latency to initiate the second hold-down, even though mice could not predict what it would be. Accordingly, a single experience of high frequency dopamine axon stimulation in the medial accumbens shell more strongly reinforces and disengages self-stimulation behavior than does stimulation at lower frequencies. Further, the capacity of augmented neurotransmission at D1 receptors and blunted neurotransmission at D2 receptors to modulate the mean duration of self-stimulation hold-downs was present on the first hold-down, confirming that these pharmacological manipulations affected the disengaging property of medial accumbens dopamine as it arose.

Whether systemic pharmacological modulation of dopamine neurotransmission influenced the duration of self-stimulation hold-downs through local actions in the medial accumbens shell, or more broadly, remains speculative. It is intriguing, however, that high frequency stimulation of dopamine inputs to the medial accumbens shell and augmenting neurotransmission at D1 receptors with a D1 agonist were both sufficient to shorten self-stimulation hold-downs and bouts of consummatory behavior. Prior studies have identified two potential circuit mechanisms that may uniquely relate to the medial accumbens shell. First, a small population of non-canonical D1 neurons in this area project to the ventral pallidum, and stimulation of this pathway induces behavioral avoidance^48^. Second, stimulation of canonical D1 neurons in the medial accumbens shell that project to the ventral tegmental area supports neither approach nor avoidance, instead inducing a general state of *behavioral suppression,* which may reflect competition between reinforcing and disengaging processes^49^. Thus, D1 neurons in the medial accumbens shell are well positioned to subserve the disengaging properties of dopamine signaling. Nevertheless, we cannot exclude the possibility that broad activation of D1 receptors serves to disengage behavior. It will be valuable for future studies to examine whether dopamine signaling in other areas like the dorsal striatum and lateral accumbens shell also promotes behavioral disengagement.

Other neurotransmitters released by dopamine axons in the medial accumbens shell, such as glutamate and γ-aminobutyric acid, may contribute to the disengaging property of this substrate. When mice were given access to five apertures that were armed with 40 Hz optogenetic stimulation of different durations (i.e., 1, 5, 20, 40 s), they nose poked more in the 5 s aperture when it triggered stimulation of ventral tegmental area dopamine neurons and more in the 1 s aperture when it triggered stimulation of neurons that co-release glutamate and dopamine^29^. However, a separate study found that stimulation of efferent ventral tegmental area glutamate fibers in the nucleus accumbens elicited a place preference only when the fibers were made unable to synthesize dopamine, which suggests there is an aversive component to the release of dopamine from this pathway. Moreover, disruption of glutamate synthesis in dopamine co-releasing afferents to the medial accumbens shell rendered stimulation of this pathway aversive^31^. Thus, dopamine release from ventral tegmental area inputs to the medial accumbens shell is critical for this pathway to drive avoidance, potentially by evoking a sensation that quickly turns aversive.

It is unclear whether the disengaging property of dopamine signaling is inherently valenced, although mice similarly reduce the duration of bouts of licking sweet solutions when they are adulterated with aversive bitter additives, which we reaffirm. We show also that neurotransmission at D1 receptors or stimulating dopamine inputs to the medial accumbens shell similarly reduce the duration of lick bouts. Related work demonstrates that manipulations of accumbens shell activity that elicit naturalistic aversive reactions like defensive treading, distress calls, or escape behavior require neurotransmission at D1 receptors, further implicating this substrate in an aversive process^50,51^. Intriguingly, a recent report confirms that high frequency stimulation of ventral tegmental area dopamine neurons evokes a distinct sensory experience, as cues paired with this stimulation selectively invigorate the pursuit of self-stimulation and not other reinforcers such as food (Pavlovian-to-instrumental transfer)^26^. Thus, dopamine activity in the medial accumbens shell may elicit a specific sensory experience that promotes rapid disengagement.

Striatal dopamine is believed to support behavior^52^ by serving as a reinforcer^53^ and prediction error signal^54^. Decades of preclinical research manipulating and observing dopamine activity during behavior support this idea, as does the indiscriminate capacity of drugs that elevate dopamine to reinforce behavior^55^. However, much of this work considers only the frequency of action initiation, rather than the duration of discrete engagements with the reinforcer. Even still, some findings hinted at a disengaging property of dopamine. For example, psychostimulants paired with food consumption^56,57^, even when administered volitionally^58^, reduced food consumption while encouraging food approach^59^, simultaneously reinforcing and disengaging behavior. Further, self-administration of psychostimulants is reduced, not amplified, by microinfusion of dopamine agonists into the accumbens^60^, and rodents prefer brief low-dose infusions of cocaine when circulating levels of cocaine in their blood, and consequently striatal dopamine levels, are high^23^. These findings point to an inherent role of dopamine receptor activation in disengaging highly reinforced actions.

Behavioral reinforcement and disengagement may be inseparably intertwined to ensure that highly reinforced actions are not maintained indefinitely. A comparable phenomenon was recently described in the dorsal striatum, where dopamine was found to not only reinforce motor actions but also increase the likelihood of uncommon action sequences^18^. This increase in action sequence variability may similarly serve to limit the duration of highly reinforced actions. Accordingly, disengagement may be a distributed property of striatal dopamine to ensure that highly reinforced actions do not become perseverative, arising from action sequence variability in the dorsal striatum and aversion in the medial accumbens shell.

By allowing mice to control not only the frequency but also the duration of self-stimulation behavior, we uncover dissociable reinforcing and disengaging properties of medial accumbens shell dopamine. Specifically, intense dopamine activity in the medial accumbens shell is strongly reinforcing but it does not support long duration lever hold-downs or lick bouts. The reinforcing and disengaging properties of medial accumbens shell dopamine are differentially affected by neurotransmission at dopamine receptors. Notably, augmenting neurotransmission at D1 receptors serves to enhance the disengaging property of medial accumbens shell dopamine. We posit that dopamine neurotransmission at D1 receptors serves to disengage the reinforcing properties of medial accumbens shell dopamine to prevent behavioral perseveration.

## STAR METHODS

### KEY RESOURCES TABLE

| REAGENT or RESOURCE | SOURCE | IDENTIFIER |
| --- | --- | --- |
| Bacterial and virus strains |  |  |
| AAV5-EF1a-DIO-hChR2(H134R)-eYFP | Canadian Neurophotonics Platform | KD3-aav5 |
| pAAV9-hSyn-GRAB-rDA1m | addgene | 140556 |
| Experimental models: Organisms/strains |  |  |
| Mouse: DAT-Cre | The Jackson Laboratory | 020080 |
| Software and algorithms |  |  |
| MED-PC V | Med Associates | N/A |
| Doric Neuroscience Studio Software | Doric Lenses | N/A |
| GuPPy | https://github.com/LernerLab/GuPPy | N/A |
| Prism v9.5.1 | Graphpad | N/A |
| SPSS v27 | IBM | N/A |
| Other |  |  |
| 473 nm DPSS Laser | LaserGlow | R471005GX |
| 300 μm core optical fiber | ThorLabs | FT300UMT |
| 200 μm core optical fiber | ThorLabs | FT200UMT |
| Nanoject II Injector | Drummond Scientific | 3-000-204 |

### RESOURCE AVAILABILITY

#### Lead contact

Further information and requests should be directed to the lea contact, Dr. Jonathan Britt.

#### Materials availability

This study did not generate new unique reagents.

#### Data and code availability

Any additional information required to reanalyze the data reported in this paper is available from the lead contact upon request. All data reported in this paper will be shared by the lead contact upon request. This paper does not report original code.

### EXPERIMENTAL MODEL AND STUDY PARTICIPANT DETAILS

#### Animals

Subjects were DAT-Cre (14f, 13m) and wild-type (n=9f, 7m) C57BL/6 experimentally naïve mice (25-30 g, 8 weeks old; bred in-house) that were single or pair-housed in standard cages (31 cm x 16.5 cm x 12 cm) under a reversed 12-hour light/dark cycle (Lights ON 9 AM, OFF 9 PM) with unrestricted access to standard chow and water. All procedures were conducted in the dark phase and in accordance with the Canadian Council on Animal Care and the McGill Animal Care Committee.

### METHOD DETAILS

#### Surgery

DAT-Cre transgenic mice were anesthetized with ketamine (Ventoquinol, 100mg/kg) and xylazine (Bayer, 10mg/kg) before being secured in a stereotaxic apparatus. For twenty Dat-Cre mice (10f, 10m) and six wild-type control mice (3f, 3m), the viral construct AAV5-EF1a-DIO-hChR2(H134R)-eYFP (Neurophotonics, 5.0×10^12^ GC/ml) was microinfused using a Nanoject II injector (Drummond Scientific, 3-000-203-G/X, 10 μm tip) in the ventral tegmental area (10° angle, AP −3.3 mm, ML ±1.14 mm, DV −4.26 to −3.96 mm; 5 min infusion, 10 min diffusion). In the same surgery optic fibers (200 μm) were implanted bilaterally in the medial accumbens shell (10° angle, AP 1.5 mm, ML ±1.52 mm, DV −4.16 mm). Seven other Dat-Cre mice (4f, 3m) had an additional viral construct, pAAV9-hSyn-GRAB-rDA1m (addgene, 5.0×10^12^ GC/ml), microinfused in the medial accumbens shell (10° angle, AP 1.5 mm, ML ±1.52 mm, DV −4.16 mm; 5 min infusion, 10 min diffusion) prior to implantation of the fiber optic cable. Mice were left to recover for 2-3 months in their home-cage allowing sufficient viral expression in dopamine neurons.

#### Apparatus

Behavioral training occurred in six conditioning chambers (ENV-307W-CT; Med-Associates Inc.) enclosed in sound-attenuating, fan-ventilated (ENV-025F) melamine cubicles (42.5 x 62.5 x 42 cm). Each chamber had a stainless-steel bar floor, paneled aluminum sidewalls, and a clear polycarbonate rear wall, ceiling, and front door. The upper left wall featured a central white house-light (ENV-215M). The right wall contained two retractable levers (ENV-312-3) located on each side of a central fluid-port equipped with two lickometer bowls (ENV-303LP2-3). All experimental events were controlled and recorded using Med PC-V software.

#### Fiber photometry recordings of optically evoked dopamine release

Dopamine cell bodies in the ventral tegmental area were photostimulated while rGRAB-DA1m fluorescence was measured from the medial accumbens shell with a 1-site, 2-color fiber photometry system from Doric Lenses. Dopamine insensitive fluorescence was assessed with an isosbestic 405 nm LED that oscillated in intensity from 10 to 100 µW at 210 Hz. Dopamine sensitive rGRAB-DA1m fluorescence was assessed with 560 nm light that oscillated in intensity from 10 to 200 µW at 450 Hz. Emitted light was measured with a 2151 Femtowatt Photoreceiver and digitized at 6 kHz with an RZ5P signal processor (Tucker-Davis Technologies). This signal was demodulated at the 210 and 450 Hz carrier frequencies and lowpass filtered at 6 Hz. The mean value of the 405 nm control signal was subtracted from the 560 nm rGRAB-DA1m signal and divided by a rolling 30s baseline over the entire session. A DPSS laser was used to deliver 10 mW 473 nm laser light to the ventral tegmental area at 1, 5, 10, 20, or 40 Hz for 1 s, or at 40 Hz for .5, 1, 3, 5, 10 s (5ms pulse) every 20s (10 trials/25 minutes). The order of stimulation frequency or duration was generated randomly.

#### Lever hold-down self-stimulation

DAT-Cre mice received 40 self-stimulation training sessions (36 min) consisting of trials (30s; 48 trials/session) initiated and terminated with the extension and retraction, respectively, of active and inactive levers. In each trial, the active lever was pseudorandomly assigned to deliver pulsed laser stimulation (5 ms; 473 nm; 10 mW) at 2.5, 10, or 40 Hz such that one third of all trials represented each frequency. Trials occurred around a variable time inter-trial interval that was drawn from an exponential distribution from 2-18 s (m=10 s). When mice performed hold-downs by depressing the active lever, laser-light was delivered through an implanted optic fiber to the medial accumbens shell, which continued so long as the lever remained depressed and terminated when the lever was released. Local stimulation of dopamine terminals in the striatum has been shown previously to evoke dopamine release^61–63^. Inactive lever hold-downs had no consequence.

After training, the contributions of neurotransmission at D1 and D2 receptors to self-stimulation of medial accumbal shell dopamine inputs were tested. Prior to each test session mice received an intraperitoneal injection of raclopride (D2 antagonist, 0.03 mg/kg), quinpirole (D2 agonist, 0.125 mg/kg), SCH23390 (D1 antagonist, 0.02 mg/kg), SKF38393 (D1 agonist, 0.25 mg/kg) or saline. Three sessions for each treatment (15 total) occurred according to a Latin-square design. Retraining sessions without injections intervened each test session to mitigate potential treatment carryover effects.

#### Consummatory behavior

DAT-Cre mice received unlimited access to a sweetened condensed milk solution (10% *v/v*) and water daily for 36 min until they made ∼200 bouts of sweetened condensed milk consumption. Bouts were initiated when mice made three licks in less than 1 s and continued until there was a 1 s pause in licking. Each lick bout has an equal probability (25%) of being paired with no stimulation (NS) and 2.5, 10, and 40 Hz optogenetic stimulation of dopamine inputs to the medial accumbens shell.

Wild-type mice received unlimited access to an 8% sucrose solution for six sessions after being treated with saline (i.p.), SKF38393 (0.125 mg/kg), or having the 8% sucrose solution adulterated with quinine (1 mM) according to a Latin-square design.

#### Histology

Mice were deeply anesthetized with sodium pentobarbital (Euthanyl^TM^, 270 mg/kg) and perfused with a 4% paraformaldehyde 0.1 M phosphate-buffered saline solution. Brains were extracted and cryoprotected in a 4% paraformaldehyde 30% sucrose solution for 1-2 days before being frozen at −80°C. Brain tissue was sliced into 50 μm coronal sections on a Lecia^TM^ cryostat, thaw-mounted on microscope slides, and cover-slipped with a MOWIOL + DAPI solution, then imaged using a Leica^TM^ epifluorescent microscope to identify viral expression in the ventral tegmental area and optical fiber tracts in the medial accumbens shell.

### QUANTIFICATION AND STATISTICAL ANALYSIS

MED-PC V software controlled and recorded the timing of all experimental events. Fiber photometry recordings were processed using GuPPy^64^. The number, duration, and latency of hold-downs and lick bouts were analyzed using repeated measures analysis of variance with the factors Treatment, Lever, and Stimulation Frequency in Prism^TM^ v9.5.1 or SPSS v27. The Kolmogorov-Smirnov test was used to compare distributions. Sessions were blocked into groups of five (training) or three (test) sessions to ensure enough hold-downs occurred to calculate mean hold-down duration. All analyses were conducted on raw data, although difference scores are shown to decompose significant interactions and visualize individual data. Corresponding figures which show absolute and normalized data is detailed in Table S1. Two mice did not perform a hold-down at one set of frequency-treatment conditions during the test phase and their data was excluded from test analyses. Post-hoc t-tests were Bonferroni-corrected (one family, all possible comparisons) and multiplicity-adjusted^65^. All analyses used α=.05.

## Acknowledgments

1. M. D. V. was supported by a postdoctoral fellowship from the Fonds de recherche du Québec – Santé (FRQS) and a Brain Star Award from the Institute of Neurosciences, Mental Health, and Addiction of the Canadian Institutes of Health Research (CIHR). N. M-L. E. was supported by a graduate fellowship from the FRQS. J. P. B. was funded by the Natural Sciences and Engineering Research Council (05100-2020), the Canadian Research Council (950-233013), and the G. W. Stairs fund at McGill University (jointly held by M. D. V. and J. P. B.). The authors thank Dr. Bruno Giros for providing some of the mice used in this study.

## Author contributions

M.D.V. and J.P.B. designed research; M.D.V., N.M-L.E., M.M.M., I.A., B.N.T., and N.M.M. performed research; M.D.V. analyzed data; M.D.V. and J.P.B. wrote the paper.

