## Supplementary Figures and Table for "A disengaging property of dopamine signaling"

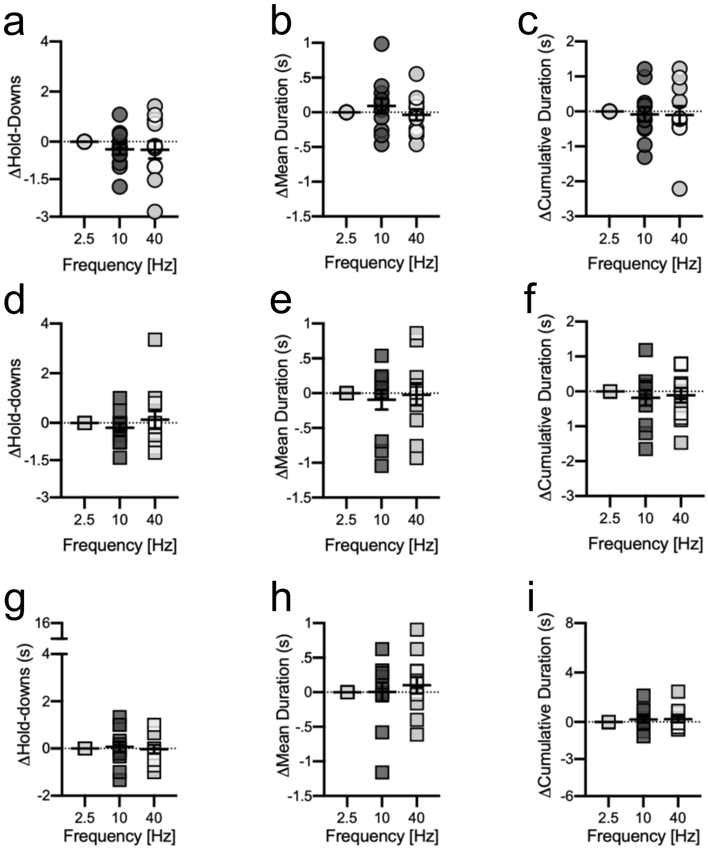


**Figure S1. Early in training self-stimulation of dopamine inputs to the medial accumbens shell is not frequency-dependent.** During the first block of training, the number of **a**, active and **d**, inactive lever hold-downs did not differ across stimulation frequencies [Lever x Frequency, $F_{(2, 22)}$=.95, $p$=.402; Frequency, $F_{(2, 22)}$=.8, $p$=.46], but more active than inactive lever hold-downs were performed [Lever, $F_{(1, 22)}$=7.12, $p$<.022]. The mean duration [Lever x Frequency, $F_{(2, 22)}$=.33, $p$=.72; Lever, $F_{(1, 22)}$=2.4, $p$<.15; Frequency, $F_{(2, 22)}$=.29, $p$=.75] and cumulative duration [Lever x Frequency, $F_{(2, 22)}$=.28, $p$=.76; Lever, $F_{(1, 22)}$=4.07, $p$<.07; Frequency, $F_{(2, 22)}$=.11, $p$=.90] of **b-c**, active and **e-f**, inactive lever hold-downs were similar across stimulation frequencies during this first block. During the last block of training the **g**, number, **h**, mean, and **i**, cumulative duration of inactive lever hold-downs did not vary across stimulation frequencies. Data are shown as within-subject differences from the lowest stimulation frequency (2.5 Hz). Averaged data are mean ± s.e.m. Symbols represent data from individual mice.

### Supplementary Figure 1

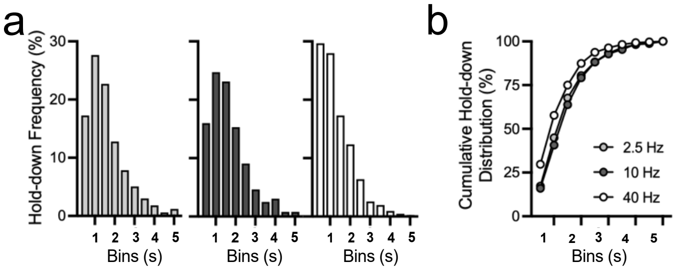


**Figure S2. Mice rapidly disengage high frequency dopamine stimulation. a**, At the end of training the distributions of lever hold-down durations during 2.5 (left), 10 (middle), and 40 (right) Hz trials were positively skewed. **b**, The cumulative distribution of hold-down durations during 40 Hz trials was significantly leftward shifted relative to 2.5 Hz and 10 Hz trials [2.5 Hz, D=.15, $p<.001$; 10 Hz, D=.18, $p<.001$] . Averaged data are mean ± s.e.m.

Supplementary Figure 2

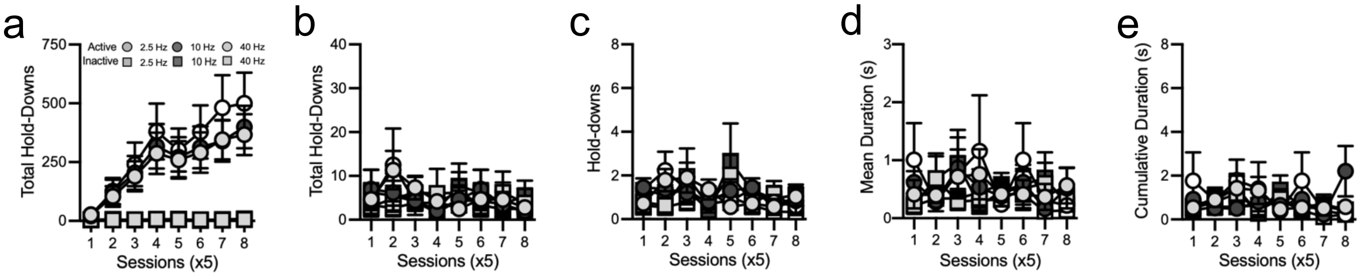


**Figure S3. Mice lacking opsins do not self-stimulate. a**, DAT-Cre mice (n=12; 7f, 5m) expressing channel rhodopsin-2 in dopamine terminals came to perform more active than inactive lever hold-downs, particularly during higher frequency trials, across training [Lever x Frequency x Session, $F_{(14,\ 154)}$=1.97, *p*=.02; Frequency x Session, $F_{(14,\ 154)}$=1.81, *p*=.04; Lever x Session, $F_{(7,\ 77)}$=8.35, *p*<.001; Lever x Frequency, $F_{(2,\ 22)}$=3.81, *p*=.038; Session, $F_{(7,\ 77)}$=8.32, *p*<.001; Frequency, $F_{(2,\ 22)}$=3.736, *p*=.04; Lever, $F_{(1,\ 11)}$=14.280, *p*=.003]. Differently, Wild-type control mice (n=6; 3f, 3m) without channel rhodopsin-2 expressed in dopamine terminals did not acquire self-stimulation behavior. **b**, In these control mice the total number of active and inactive hold-downs were seldom and similar [Lever x Frequency x Session, $F_{(14,\ 56)}$=.47, *p*=.94; Frequency x Session, $F_{(14,\ 56)}$=1.24, *p*=.28; Lever x Session, $F_{(7,\ 28)}$=1.72, *p*=.15; Lever x Frequency, $F_{(2,\ 8)}$=.6, *p*=.57; Session, $F_{(7,\ 28)}$=.78, *p*=.61; Frequency, $F_{(2,\ 8)}$=2.17, *p*=.18; Lever, $F_{(1,\ 4)}$=.001, *p*=.98]. **c**, Consistently, a similar number of active and inactive hold-downs per trial were performed which showed some spurious changes across frequencies [Lever x Frequency x Session, $F_{(14,\ 56)}$=1.36, *p*=.21; Frequency x Session, $F_{(14,\ 56)}$=2.2, *p*=.019; Lever x Session, $F_{(7,\ 28)}$=1.78, *p*=.18; Lever x Frequency, $F_{(2,\ 8)}$=.95, *p*=.43; Session, $F_{(7,\ 28)}$=1.27, *p*=.3; Frequency, $F_{(2,\ 8)}$=.96, *p*=.43; Lever, $F_{(1,\ 4)}$=.15, *p*=.72]. **d**, The average duration of active and inactive hold-downs were similarly brief [Lever x Frequency x Session, $F_{(14,\ 56)}$=.97, *p*=.5; Frequency x Session, $F_{(14,\ 56)}$=.91, *p*=.56; Lever x Session, $F_{(7,\ 28)}$=1.22, *p*=.32; Lever x Frequency, $F_{(2,\ 8)}$=.88, *p*=.45; Session, $F_{(7,\ 28)}$=.95, *p*=.48; Frequency, $F_{(2,\ 8)}$=.28, *p*=.76; Lever, $F_{(1,\ 4)}$=.32, *p*=.61]. **e**, Cumulatively, control mice spent little time holding down the active and inactive levers [Lever x Frequency x Session, $F_{(14,\ 56)}$=1.39, *p*=.19; Frequency x Session, $F_{(14,\ 56)}$=1.42, *p*=.18; Lever x Session, $F_{(7,\ 28)}$=1.49, *p*=.21; Lever x Frequency, $F_{(2,\ 8)}$=.83, *p*=.47; Session, $F_{(7,\ 28)}$=1.18, *p*=.34; Frequency, $F_{(2,\ 8)}$=.11, *p*=.89; Lever, $F_{(1,\ 4)}$=.43, *p*=.55]. Data from one control mouse was only collected for 30 sessions and was excluded from analysis. Averaged data are mean ± s.e.m.

Supplementary Figure 3

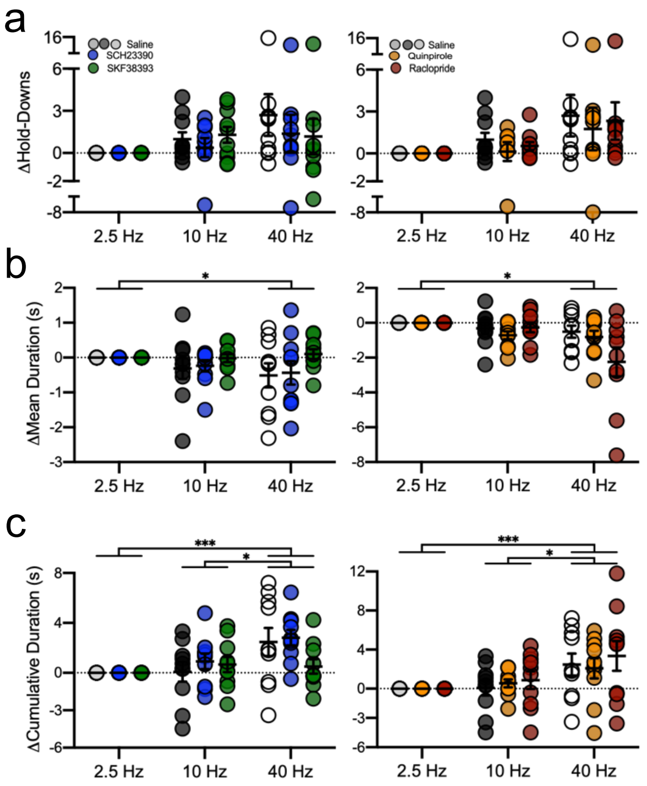


**Figure S4. During drug treatment sessions higher stimulation frequencies reduced lever hold-down durations while increasing cumulative hold-down times. a,** The number of active lever hold-downs per trial did not vary significantly across trials [Frequency, $F_{(2, 18)}$=1.77, *p*=.198]. **b,** The mean duration of active lever hold-downs was briefer during 40 Hz trials than 2.5 Hz trials [$t_{(9)}$=3.36, *p*=.011; Frequency, $F_{(2, 18)}$=5.72, *p*=.012]. **c,** Mice performed longer cumulative active lever hold-downs during 40 Hz trials than 10 Hz and 2.5 Hz trials [10 Hz, $t_{(9)}$=3.31, *p*=.01; 2.5 Hz, $t_{(9)}$=4.54, *p*<.001; Frequency, $F_{(2, 18)}$=11.04, *p*<.001]. Data are shown as within-subject differences from 2.5 Hz trials. Averaged data are mean ± s.e.m. Symbols represent data from individual mice. Asterisks indicate Bonferroni-corrected and multiplicity-adjusted post-hoc comparisons, *p*<.001***, *p*<.05*.

### Supplementary Figure 4

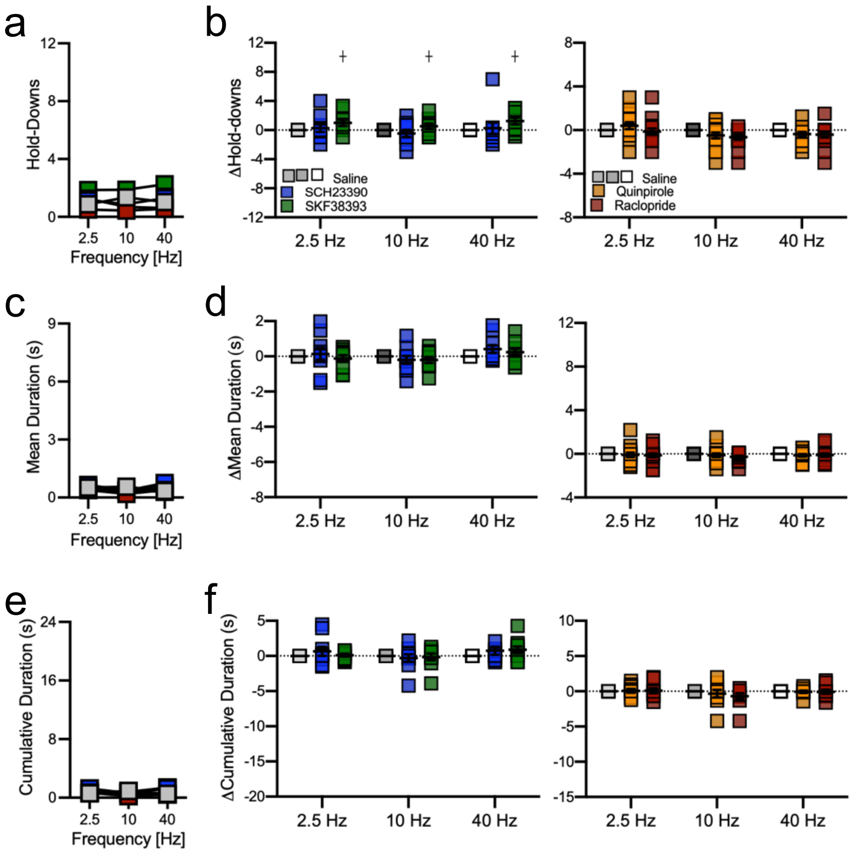


**Figure S5. The cumulative and mean duration of inactive hold-downs were unaffected by dopamine receptor agonists and antagonists.** While the **a**, number of inactive lever hold downs was unaffected by stimulation frequency [Frequency x Treatment, $F_{(8, 72)}$=.57, $p$=.8; Frequency, $F_{(2, 18)}$=.19, $p$=.83], there was a significant effect of treatment [Treatment, $F_{(4, 36)}$=9.81, $p$<.001]. **b**, The D1 agonist SKF38393 elevated the number of inactive hold-downs [$t_{(9)}$=3.66, $p$=.008], whereas other treatments had no effect [SCH23390, $t_{(9)}$=.09, $p$>.99; Quinpirole, $t_{(9)}$=.92, $p$>.99; Raclopride, $t_{(9)}$=2.32, $p$=.26]. The **c-d**, mean duration [Frequency x Treatment, $F_{(8, 72)}$=1.11, $p$=.37; Treatment, $F_{(4, 36)}$=.86, $p$=.5; Frequency, $F_{(2, 18)}$=1.72, $p$=.211] and **e-f**, cumulative duration [Frequency x Treatment, $F_{(8, 72)}$=.84, $p$=.57; Treatment, $F_{(4, 36)}$=2.14, $p$=.1; Frequency, $F_{(2, 18)}$=1.37, $p$=.28] of inactive lever hold-downs were wholly unaffected by drug treatments and (active lever) stimulation frequency. Data in panels *b*, *d*, and *f* are shown as within-subject differences from the saline treatment condition. Averaged data are mean ± s.e.m. Symbols represent data from individual mice. Crosses indicate Bonferroni-corrected and multiplicity-adjusted simple effects comparisons of treatment relative to saline $p$<.0001┼.

Supplementary Figure 5

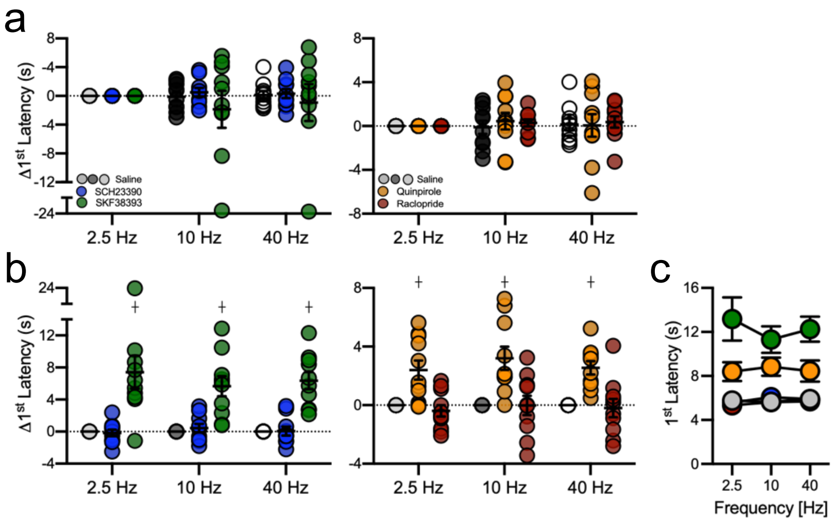


**Figure S6. The latency of the first self-stimulation lever hold-down. a**, The latency to initiate the first active lever hold-down after trial onset did not vary across stimulation frequencies [Frequency, $F_{(2, 18)}$=.049, *p*=.95]; however, **b**, mice were slower following treatments [Treatment, $F_{(4, 36)}$=40.94, *p*<.001] with the D1 agonist SKF38393 [$t_{(9)}$=10.18, *p*<.001] and the D2 agonist quinpirole [$t_{(9)}$=4.40, *p*<.001]. The D1 antagonist SCH23390 [$t_{(9)}$=.13, *p*>.99] and the D2 antagonist raclopride [$t_{(9)}$=.34, *p*>.99] had no effect. **c**, Summary of the unnormalized data used for analysis of the mean latency to perform the first hold-down. Data are shown as within-subject differences from the 2.5 Hz trials in *a* and from the saline treatment condition in *b*. Averaged data are mean ± s.e.m. Symbols represent data from individual mice. Crosses indicate Bonferroni-corrected and multiplicity-adjusted simple effects comparisons of treatment relative to saline *p*<.0001┼.

Supplementary Figure 6

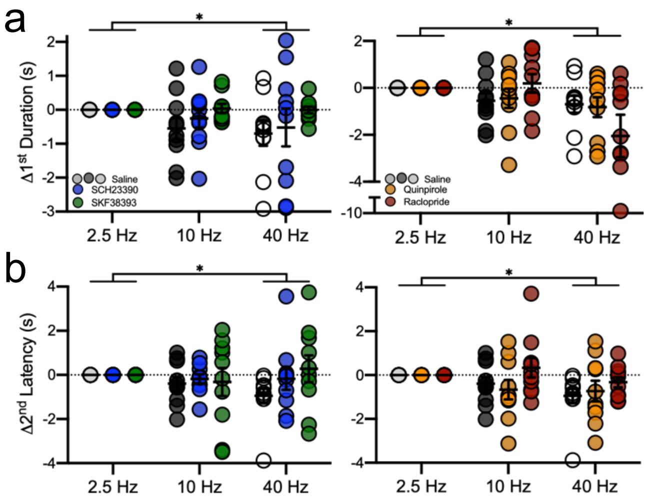


**Figure S7. The duration of the first and latency to initiate the second active lever hold-down are stimulation frequency dependent. a**, The mean duration of the first active lever hold-down of each trial varied with stimulation frequency [Frequency, $F_{(2,\ 18)}$=5.01, $p$=.019] and was briefer during 40 Hz trials relative to 2.5 Hz trials [$t_{(9)}$=3.04, $p$=.02]. **b**, The latency to initiate the second active lever hold-down, after termination of the first, varied across stimulation frequencies [Frequency, $F_{(2,\ 18)}$=4.02, $p$=.04] and was shorter during 40 Hz trials than 2.5 Hz trials [$t_{(9)}$=2.79, $p$=.04]. Data are shown as within-subject differences from 2.5 Hz trials. Averaged data are mean ± s.e.m. Symbols represent data from individual mice. Asterisks indicate Bonferroni-corrected and multiplicity-adjusted post-hoc comparisons, $p$<.05*.

Supplementary Figure 7

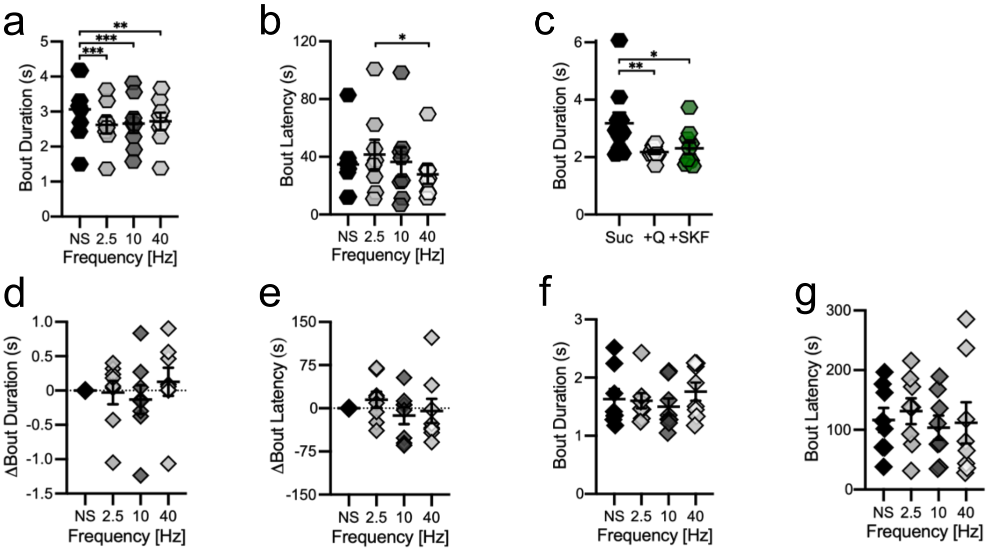


**Figure S8. Dopamine axon stimulation and D1 receptor activation reduce lick bout durations.** Summary of unnormalized data used for analysis of **a**, bout duration, **b**, latency to a subsequent bout of sweetened condensed milk consumption, and **c**, bout duration during sucrose solution consumption. Water was concurrently available in both experiments, but bouts were too few to warrant analysis during the sucrose experiment. When mice received access to sweetened condensed milk paired with optogenetic stimulation of dopamine inputs to the medial accumbens shell stimulation also occurred on a subset of water bouts which may have increased the frequency of water bouts. Optogenetic stimulation did not affect the **d**,**f**, duration or **e**,**g**, latency of water bouts, although these bouts were short and seldom [Duration: Frequency, $F_{(3, 21)}$=.74, $p$=.53; Latency: Frequency, $F_{(3, 21)}$=.48, $p$=.697]. Data in *d* and *e* are shown as within-subject differences from the no stimulation condition (NS). Averaged data are mean ± s.e.m. Symbols represent data from individual mice. Asterisks indicate Bonferroni-corrected and multiplicity-adjusted post-hoc comparisons, $p$<.001***, $p$<.01**, $p$<.05*.

Supplementary Figure 8

| Exp. | n | Figure(s) and data normalization | Description |
|---|---|---|---|
| 1 | 7 | Baseline normalized: 1e-f | Photostimulation of ventral tegmental area dopamine neurons produces frequency- and duration-dependent dopamine release in the medial accumbens shell. |
| 2a | 12 | Absolute: 1g-i (left), S3a<br>Frequency normalized: 1g-i (right), S1<br>Other: S2 | Acquiring self-stimulation of dopamine inputs to the medial accumbens shell. |
| 2b | 10 | Absolute: 2a,c,e; S5a,c,e<br>Frequency normalized: S4<br>Vehicle normalized: 2b,d,f; S5b,d,f | Frequency dependent self-stimulation is modulated by dopamine receptor agonists and antagonists. |
| 2c | 10 | Absolute: S6e<br>Frequency normalized: S6a<br>Vehicle normalized: S8b | Latency to perform the first self-stimulation hold-down of a trial is uniform across trial types. |
| 2d | 10 | Absolute: 3a<br>Frequency normalized: S7a<br>Vehicle normalized: 3b | Duration of the first self-stimulation hold-down of a trial is frequency- and treatment-dependent. |
| 2e | 10 | Absolute: 3c<br>Frequency normalized: S7b<br>Vehicle normalized: 3d | Latency to perform a second self-stimulation hold-down within a trial is frequency-dependent. |
| 3 | 6 | Absolute: S3b-e | Mice lacking opsins to not acquire self-stimulation behavior. |
| 4 | 8 | Absolute: S8a-b, S8f-g<br>Frequency normalized: 4a-b, S8d-e | Photostimulation of dopamine inputs to the medial accumbens shell disengages consummatory behavior. |
| 5 | 10 | Absolute: S8c, S8f-g<br>Vehicle normalized: Fig4c | Adulteration of a sucrose solution with quinine or treatment with SKF38393 shortens lick bout durations. |

Supplementary Table 1
